# Inflammatory Crosstalk Between Rotator Cuff Tissues is Altered with Age and Sex

**DOI:** 10.1101/2025.06.10.658924

**Authors:** Hana Kalčo, Paola Divieti Pajevic, LaDora V. Thompson, Brianne K. Connizzo

## Abstract

Musculoskeletal disorders, particularly those affecting the shoulder, are a significant health concern, especially in aging populations. Nevertheless, the initiating factors of joint degeneration remain poorly understood. Research has primarily focused on age-related changes in individual musculoskeletal tissues, with limited investigation into the complex interactions between tissues. Recent studies on interorgan communication between musculoskeletal tissues and other organs have gained attention, but local interactions within the shoulder remain underexplored. This study aims to investigate age- and sex-related differences in bone-tendon-muscle (BTM) crosstalk, hypothesizing that these interactions vary by age and sex, with older and female tissues exhibiting a reduced secretory phenotype. Using novel *in vitro* monoculture and co-cultures of explanted whole tissues, we assessed inflammatory responses across bone, tendon, and muscle from young and aged male and female C57BL/6J mice. Our results demonstrate significant age- and sex-dependent differences in cytokine secretion, with aged males and females showing altered inflammatory profiles. We observed a general increase in pro-inflammatory cytokine secretion in monocultures, with aging amplifying this response. Tissue co-cultures revealed that crosstalk between bone and tendon was primarily mediated through secreted factors, while muscle-tendon communication required physical proximity or contact, suggesting a distinct mode of interaction between these tissues. Sex differences were evident in both the individual tissue responses and in the patterns of inter-tissue communication. Importantly, our findings suggest that tendon plays a crucial role in mediating inter-tissue communication, with aging disrupting this crosstalk. However, these sex differences diminished with aging, indicating that the age-related decline in tissue-specific signaling may override sex-based distinctions.

## Introduction

Age-related joint degeneration is a significant socioeconomic and emotional burden, especially for an increasingly aged population.^1^ There are currently no available disease-modifying drugs, and often the only treatment is to replace the entire joint with artificial components. Despite the high prevalence, the initiating factors and key contributors to whole joint degeneration remain poorly understood.^2^ While extensive research has focused on age-related pathological changes in individual musculoskeletal tissues, fewer studies have examined the complex interactions between these tissues. For example, osteoarthritis (OA) was historically considered a cartilage-specific disease; however, research over the past decade has revealed that several other tissues contribute to the pathogenesis of the disease, most notably the synovium.^3^ In addition, studies exploring crosstalk between musculoskeletal tissues and other organ systems, like the kidney, brain, and gut, are gaining significant attention and funding support, especially within the context of exercise physiology. However, our understanding of interorgan communication between tissues <u>within</u> a synovial joint is still quite limited, despite the proximity, and in some cases physical connection, between them. Although they serve as a mechanical conduit between different musculoskeletal tissues, very few studies have explored the role of tendons and ligaments in mediating joint crosstalk.

Historically, tendon has often been viewed as a passive mechanical link rather than an active biological participant in joint homeostasis.^4^ However, tendon has been shown to contribute to the inflammatory environment mediated by cytokines and various immune cells in tendinopathy.^5^ We previously developed a novel *in vitro* bone-tendon-muscle (BTM) organ culture system that allows for controlled investigation of local tissue interactions. We observed that the tendon responds rapidly to a combined muscle and bone injury, demonstrating inflammation, cell death, and matrix degeneration.^6^ This work demonstrated for the first time that tendon was a biological participant in the synovial environment, and that tendon is responsive to cues from both muscle and bone. However, tendon’s response was initiated by injuries in both muscle and bone, and therefore the specific tissue driving these changes is still undetermined. Further, it has also not yet been established whether muscle and bone are also responsive to signals derived from tendon. Nevertheless, we believe that communication between tissues within the joint is likely critical to overall joint health.

Given differences in the reported rates of joint degeneration based on both sex and age, it’s also important to consider whether crosstalk might be dependent on these biological factors. Various age-related diseases have been shown to alter the secretory phenotype of both bone and muscle cells, suggesting that circulatory crosstalk is altered.^7^ Further, it’s known that the circulating inflammatory milieu becomes dysregulated over time,^4^ often referred to as the theory of “inflammaging.” Sustained elevated levels of pro-inflammatory cytokines in the joint are also implicated in the progression of osteoarthritis and other musculoskeletal disorders.^3^ However, it’s not clear how local crosstalk changes with aging or how that contributes to joint progression. With respect to sex, there is substantial evidence that joint disease progression and tissue remodeling mechanisms are different between male and female subjects.^8^ Additionally, there is likely a significant interaction between age and sex based on the high rates of joint disease among post- menopausal women. Despite this, there are no studies to date focusing on sex differences in local intertissue communication.

Therefore, the overall objective of this study is to define age- and sex-related differences in bidirectional crosstalk between individual musculoskeletal tissues, specifically bone, tendon, and muscle. We hypothesized that older and female tissues would exhibit a reduced secretory phenotype based on previous findings. By systematically evaluating the biochemical signaling between tendon and muscle, as well as tendon and bone, we seek to elucidate the underlying mechanisms contributing to musculoskeletal degeneration. This study provides a critical step toward understanding how tissue-specific aging influences whole-joint pathology, with implications for the development of targeted therapeutic interventions.

## Methods

### 2.1 Sample preparation

Explants were harvested from young (4 months old) and aged (24 months old) male and female C57BL/6J mice. Aged mice were obtained from the NIH/NIA Aging Rodent Colony, and young mice were obtained from Jackson Laboratories. We collected bone-tendon-muscle (BTM) explants from the mouse shoulder as previously described.^9^ The muscle was dissected cleanly from the supraspinous fossa to collect as much intact supraspinatus muscle as possible. We marked the borders of the supraspinatus tendon to isolate the tendon only from other surrounding tissues, and then sharply dissected the other rotator cuff tendons from the humeral head at their entheses. Bone was sharply cut at the base of the humeral head, leaving a bony piece containing both cortical and trabecular regions. Bone-only (BO) and muscle-only (MO) explants were further isolated from BTMs by transection at the enthesis and myotendinous junction of the supraspinatus tendon, respectively (Fig 1). We also isolated combined muscle and tendon co-culture (MT) and bone-tendon co-culture (BT) groups, preserving native junctions between the two tissues and only removing the third tissue (bone or muscle, respectively) (Fig 1). Finally, we also collected flexor tendon explants (FDLs) for this study to serve as our ‘tendon’ group rather than the supraspinatus tendon given their larger size, increased ease of visualization, and established analysis protocols.

**Figure 1:**
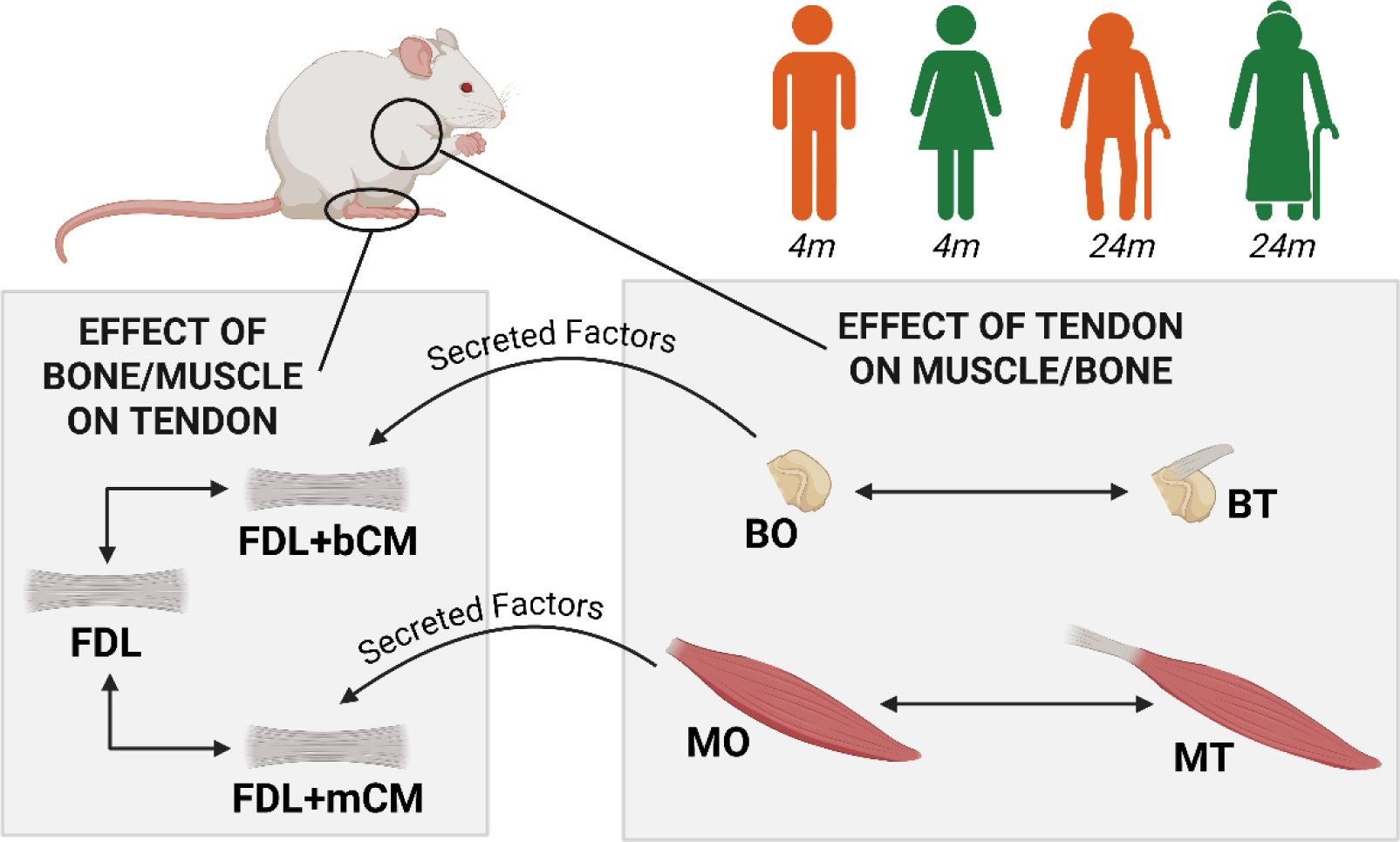
Explants were harvested from young (4 months old) and aged (24 months old) male and female C57BL/6J mice. We induced trauma via sharp dissection and isolated bone only (BO) and muscle only (MO) monocultures to collect tissue-specific signaling. Conditioned medium containing secreted factors from bone injury (bCM) and muscle injury (mCM) was then added to healthy flexor tendon explants for 24h to analyze effects of injury on tendon health. We also collected baseline tissues FDLs to address any concerns over the effect of culture conditions alone. To study crosstalk, we added combined bone-tendon co-culture (BT) and muscle-tendon co-culture (MT) groups, preserving native junctions between the two tissues. Data from BT and MT groups was normalized and compared statistically to BO and MO groups, respectively, to determine the **effect of tendon inclusion on bone or muscle responses**. Data from FDLs treated with mCM or bCM were normalized and compared statistically to FDL alone to determine the **effect of muscle or bone on tendon**.

### 2.2 Culture Conditions

After the harvest, all explants were washed in 1x PBS containing 100 units/mL penicillin G and 100 μg/mL streptomycin (Fisher Scientific), Amphotericin B, and transferred to a culture medium. The culture medium consisted of low glucose Dulbecco’s Modified Eagle Media (DMEM) (1 g/L (Fisher Scientific)) with 10% fetal bovine serum (Cytiva), and antibiotics (see above). All explants were maintained under stress deprived conditions, where they were freely suspended in the medium without mechanical loading. To study secreted factors and their effects on tendon, we collected spent medium from muscle-only and bone-only cultures after 24h and then mixed with fresh medium in a 1:1 ratio to create muscle conditioned medium (mCM) and bone conditioned medium (bCM) to be added to tendon explants.

### 2.3 Analysis of Secreted Proteins

Medium from all groups was collected to assess production of pro- and anti-inflammatory cytokines after 24h using a custom Meso Scale Diagnostics multi-spot assay, which quantifies protein levels of IFN-γ, IL-1β, IL-4, IL-6, IL-10, IL-15, KC/GRO (mouse analog of IL-8), MCP-1 and TNF-α.

### 2.7 Statistical Analyses

Data from BT and MT groups was normalized and compared statistically to BO and MO groups, respectively, to determine the effect of tendon inclusion on injury responses. Data from FDLs treated with mCM or bCM were normalized and compared statistically to FDL alone to determine the effect of muscle or bone injury, respectively. All data is presented as a fold-change relative to the control condition (tissue alone), such that a value of ‘1’ would represent no crosstalk and a value greater than one represents amplification of tissue responses. Statistical comparisons were made using unpaired two-tailed t-tests relative to control groups for each protein, and between young and aged tissues with significance set at *p<0.05.

## Results

We first looked at age-related differences in the amount of protein secreted by monocultures alone in all groups. In bone-only cultures, aged males secreted more IL- 10, IL-1β and IL-4 than young males (Fig. 2A). In tendon-only cultures, aged males secreted more IFN-γ and IL-1β (Fig. 2B). In muscle-only cultures, aged males produced more IL-6, KC/GRO and TNF-α (Fig. 2C). In bone-only cultures, aged female tendons secreted <u>less</u> MCP-1 and TNF-α in comparison to young tissues (Fig. 2D). In tendon-only cultures, aged female tendons produced more IL-6, KC/GRO and TNF-α (Fig. 2E). In muscle-only cultures, there were generally no differences between the cytokine production between young and aged female tendons, except for an increase in IFN-γ secretion with aging (Fig. 2F).

**Figure 2:**
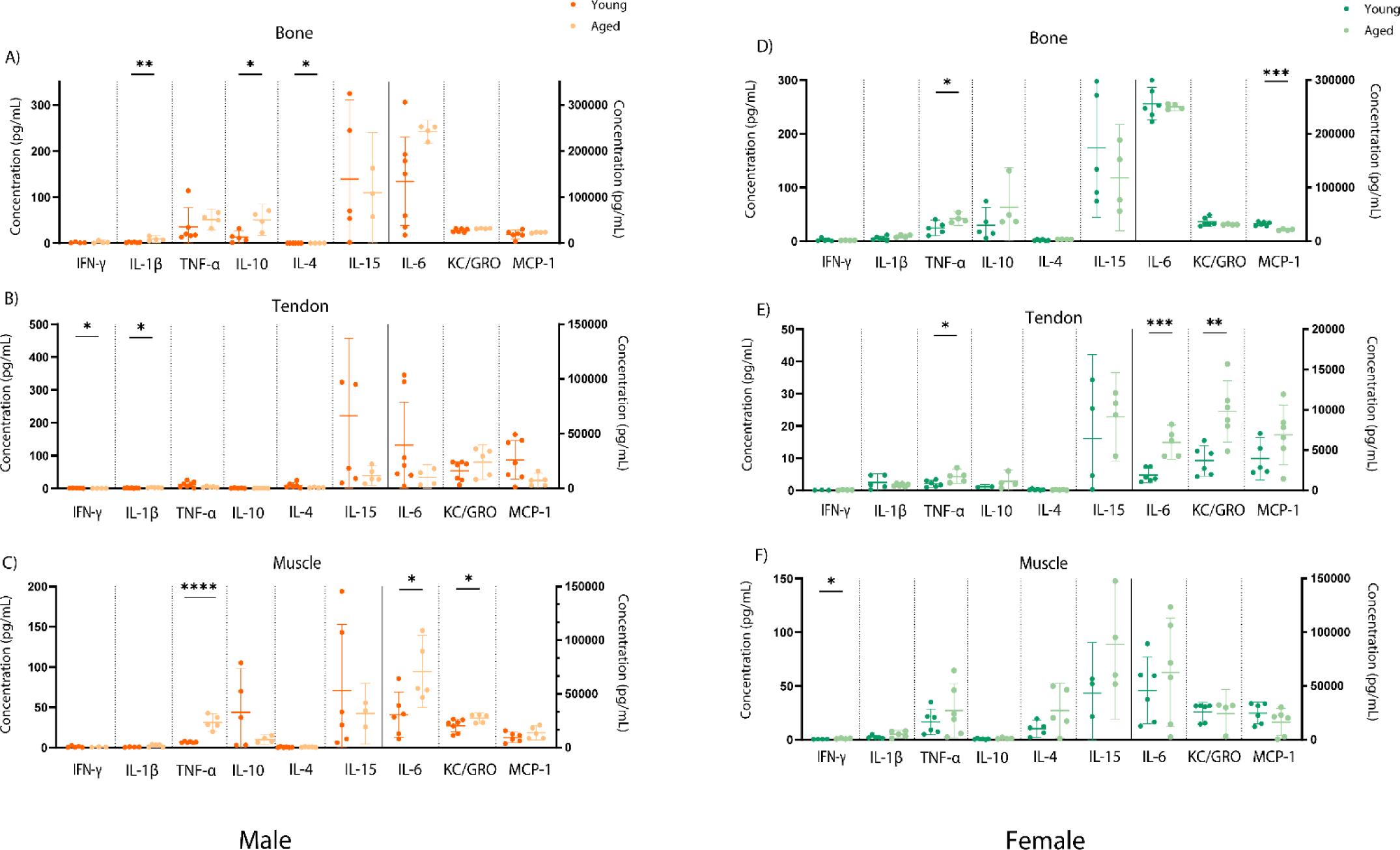
(A&D) Bone-only, (B&E) tendon-only, and (C&F) muscle-only explants were cultured independently, and secreted protein levels were measured from conditioned media to assess the inflammatory profile of each tissue type. Data represents mean (a) Bone, (b) tendon, (c) muscle only explants protein secretions in the culture. Data shown as mean ± SD with statistical significance (*p≤0.05). Due to significant differences in secreted protein levels, IL-6, KC/GRO and MCP-1 are plotted on the right axis (ng/mL) with all other proteins on the left axis (pg/mL).

We then investigated sex differences in crosstalk between muscle/bone and tendon via secreted protein interactions (Fig. 3A-B) and physical connections through co-cultures (Fig. 3C-D). In response to secreted bone factors, young female flexor tendons secreted fewer proteins overall, with significant differences in IFN-γ, IL-10, IL-15, and IL-1β (Fig.3A). However, KC/GRO and MCP-1 were found to be increased in females compared to males. The sex-related changes were fewer in response to muscle factors with significant decreases in IL-6, KC/GRO, MCP-1 and TNF-α (Fig. 3B). In both bone cultures and muscle cultures including tendon (BT, MT, respectively), we saw fewer protein secretions in females than in males. In BT (Fig. 3C), IL-10, IL-15, IL-1β, IL-4, IL-6 and MCP-1 were detected at a higher concentration in young males than young females. In MT, we saw reduced protein secretions of IFN-γ, IL-15, IL-b, IL-4, IL-6, KC/GRPO, MCP- 1 and TNF-α in young females. Interestingly, we saw no sex-related differences in the protein secretions in aged cultures (Supplemental Fig. S3).

**Figure 3:**
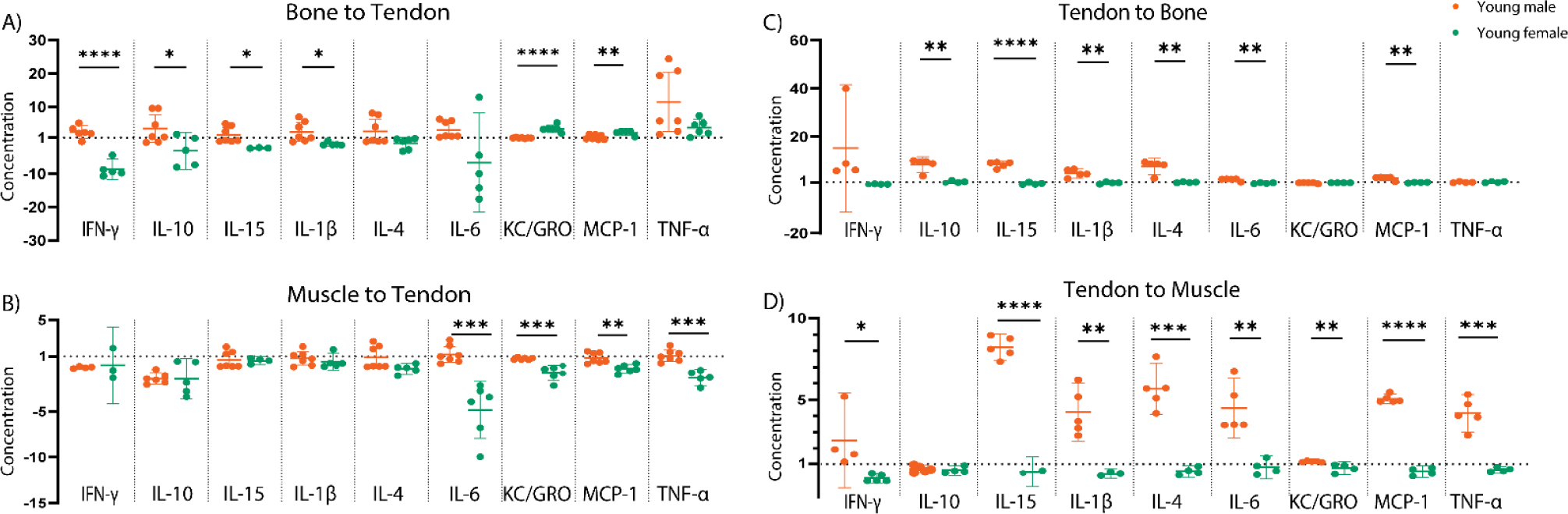
Concentration of pro- and anti- inflammatory cytokines released in spent media from young male and female CM treated FDLs (A and B) and BT and MT co-culture (C and D). All data points are normalized to bone/tendon/muscle alone. Data shown as mean ± SD. Statistical significance (*p≤0.05) between normalized young male and female represented by solid line spanning between groups.

There were also significant effects of aging on crosstalk between tendon and muscle/bone in almost all conditions. We first started with age-related differences in males. In response to bone secreted factors, aged male FDL tendons secreted less IFN- γ, IL-10, IL-6, and TNF-α (Fig. 4A). In response to secreted factors from muscle, there was an increased production of IL-10 only in aged male tissues (Fig. 4B). Similar to secreted factors, when bone and tendon were maintained in co-culture, both pro-inflammatory (IL-6, IL-1β, MCP-1) and anti-inflammatory crosstalk (IL-4, IL-10, IL-15) was significantly decreased in aged male tissues compared to young (Fig. 4C). This was seen in muscle-tendon crosstalk as well, although significance was only detected in IL-6, IL-15 and MCP-1 crosstalk (Fig. 4D).

**Figure 4:**
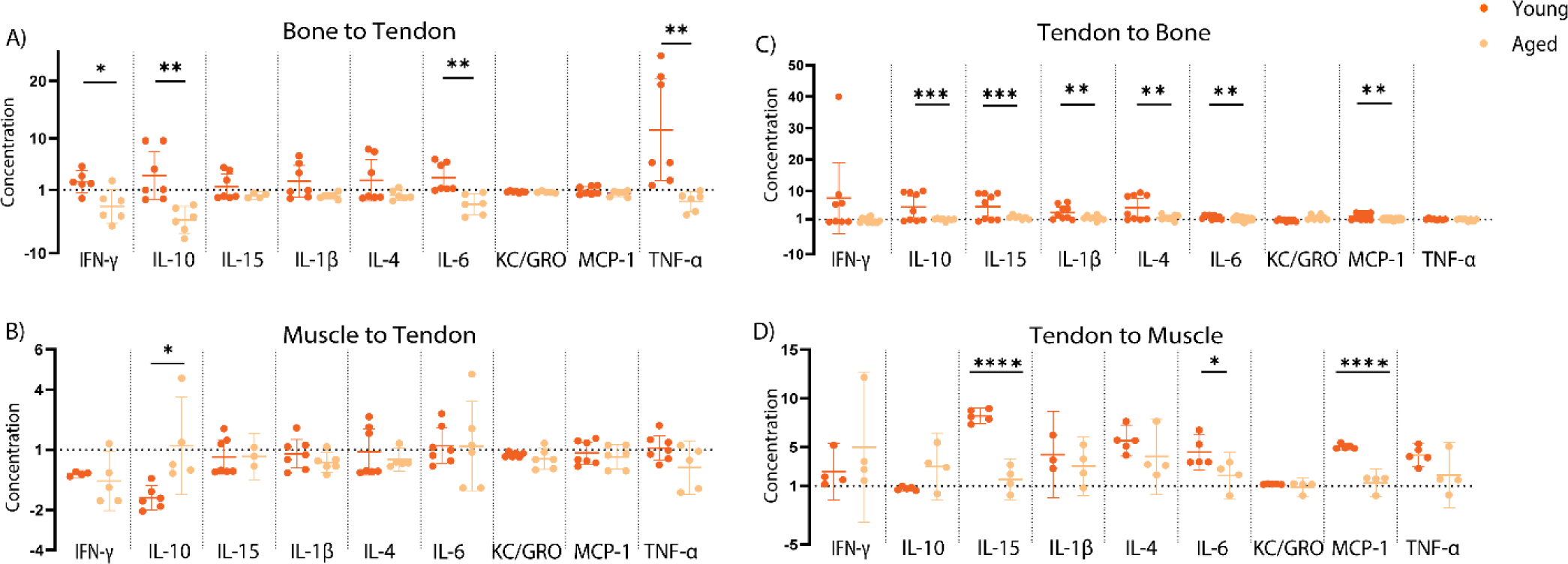
Concentration of pro- and anti- inflammatory cytokines released in spent media from young and aged male CM treated FDLs (A and B) and BT and MT coculture (C and D). All data points are normalized to bone/tendon/muscle alone. Data shown as mean ± SD. Statistical significance (*p≤0.05) between normalized young and aged represented by solid line spanning between groups.

In females, inflammatory crosstalk was generally unchanged or increased in aged tissues compared to young. In tendons treated with bone secreted factors, TNF-α and IL-15 crosstalk was increased in aged female tissues (Fig. 5A). In female tendons treated with muscle-derived factors, IL-6, TNF-α and IL-10 crosstalk was increased with aging while IL-4 and IL-15 crosstalk was decreased (Fig. 5B). In co-cultures containing female bone and tendon, IL-1β and IL-15 crosstalk was increased with aging (Fig. 5C), but interestingly there were no age-related differences in muscle-tendon crosstalk (Fig. 5D).

**Figure 5:**
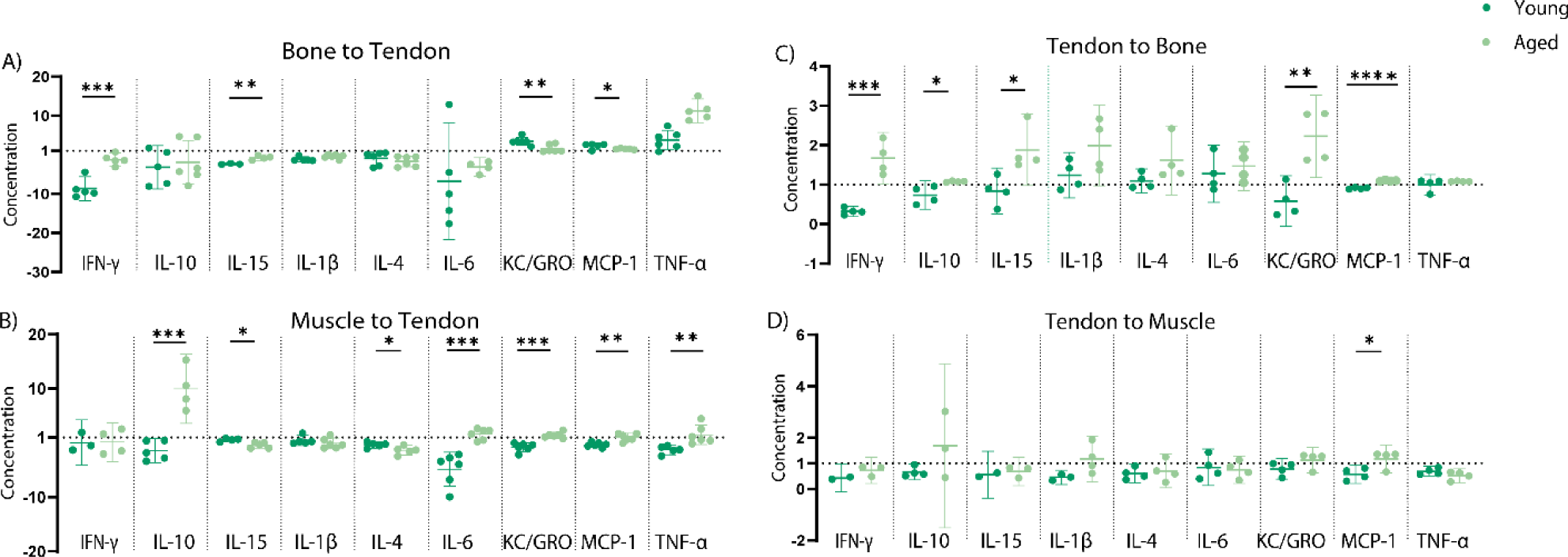
Concentration of pro- and anti- inflammatory cytokines released in spent media from young and aged female CM treated FDLs (A and B) and BT and MT coculture (C and D). All data points are normalized to bone/tendon/muscle alone. Data shown as mean ± SD. Statistical significance (*p≤0.05) between normalized young and aged represented by solid line spanning between groups.

## Discussion

This study had two main objectives: (1) to examine the independent effects of muscle- and bone-derived secreted factors and (2) to investigate the bidirectional crosstalk between tendon and adjacent injured muscle and bone. We found that tissues maintained in monocultures, regardless of sex, do exhibit secretion of several inflammatory cytokines (Fig. 2). Most notably, levels of IL-6, KC/GRO, and MCP-1 were secreted in concentrations on par with what’s previously been measured in the synovial fluid extracted from tendinopathy and arthritis patients (ng/mL).^10,10^ We believe this is due to the harvest process itself as tissues are removed via sharp dissection. Importantly, levels of cytokines secreted were higher in bone and muscle cultures (about 9-fold higher on average) than tendon cultures. This is unsurprising given the significant size and cellularity differences between tendon and the other two tissues (Fig. S1,2).

In general, the secretion of pro-inflammatory cytokines was mostly amplified with aging. This was true in all three tissues derived from male mice, as well as in tendons derived from female mice. This trend may be attributed to “inflammaging,” given the overall rise in pro-inflammatory cytokine secretion. Notably, aged male bone monocultures also exhibited elevated levels of IL-10 and IL-4, two cytokines known for their anti-inflammatory properties.^12^ This finding may be significant, as some in vivo studies have shown that daily local IL-4 injections lead to significantly decreased bone loss in mice and an increased M1/M2 ratio while IL-10 is known to increase osteogenesis.^12^ This could be compensation for age-related changes, where the upregulation of anti-inflammatory cytokines such as IL-4 and IL-10 serves to counterbalance the heightened pro-inflammatory environment associated with aging. Female bone cultures did not demonstrate this increase in IL-10 and IL-4 secretion, and overall, female bone cultures secreted fewer cytokines with aging, including MCP-1 and TNF-α. Even more strikingly, muscle monocultures derived from females demonstrated no age-related differences in the secretion of inflammatory cytokines. This is different from what we would expect to see based on previous human data.^13^ The decrease in endogenous estrogen levels during menopause is typically associated with more inflammation.^13^ However, the effects of hormonal changes with age are most likely underestimated here, as mice maintain relatively stable estrogen levels throughout aging compared to the sharp decline observed in postmenopausal humans.^14^

Moving beyond monocultures, we then investigated how bone-tendon and muscle-tendon crosstalk was altered with aging and between sexes. In general, inclusion of additional tissues, either through secreted factors in conditioned medium or by physical connections in co-culture, resulted in altered secretion of many inflammatory cytokines. Interestingly, the effect of tendon on muscle and bone in young male-derived tissues was almost always amplifying the inflammatory response, as noted by positive values in the right panels of Figures 3-4. Importantly, these changes were significantly greater than those that would be produced by the tendon alone. Considering the relative size and cellularity of the tendon (Fig. S1,2), this implies an important role of tendon as a mediator of inter-tissue communication. However, this was not true for young females or either aged group and crosstalk appeared to be quite dependent on cytokine and tissue-type. Further, responses of tendon to muscle- and bone-derived factors were sometimes inhibited in female and aged tendons, resulting in less cytokine secretion and suggesting potential feedback loops in communication.

Sex differences in intertissue crosstalk were clearly identifiable in cultures derived from young mice (Fig. 2, 6). These differences imply that biological sex plays a critical role in musculoskeletal signaling, potentially influencing injury response, healing, and long-term tissue homeostasis. All cytokines, both pro- and anti-inflammatory, seem to decrease in young females compared to young males, except for KC/GRO and MCP-1. This could be due to the immune system which functions in a sexually dimorphic manner, although typically females exhibit more robust immune responses than males.^15,16^ This is opposite from what we see in our data. Perhaps, this could be due to the absence of hormones included in the present studies, which are known to affect both immune and ECM repair responses; this will be addressed in our future studies. If young male and female tissues exhibit distinct crosstalk patterns, it implies that therapeutic approaches for musculoskeletal injuries may need to be sex-specific to best optimize healing outcomes.

**Figure 6:**
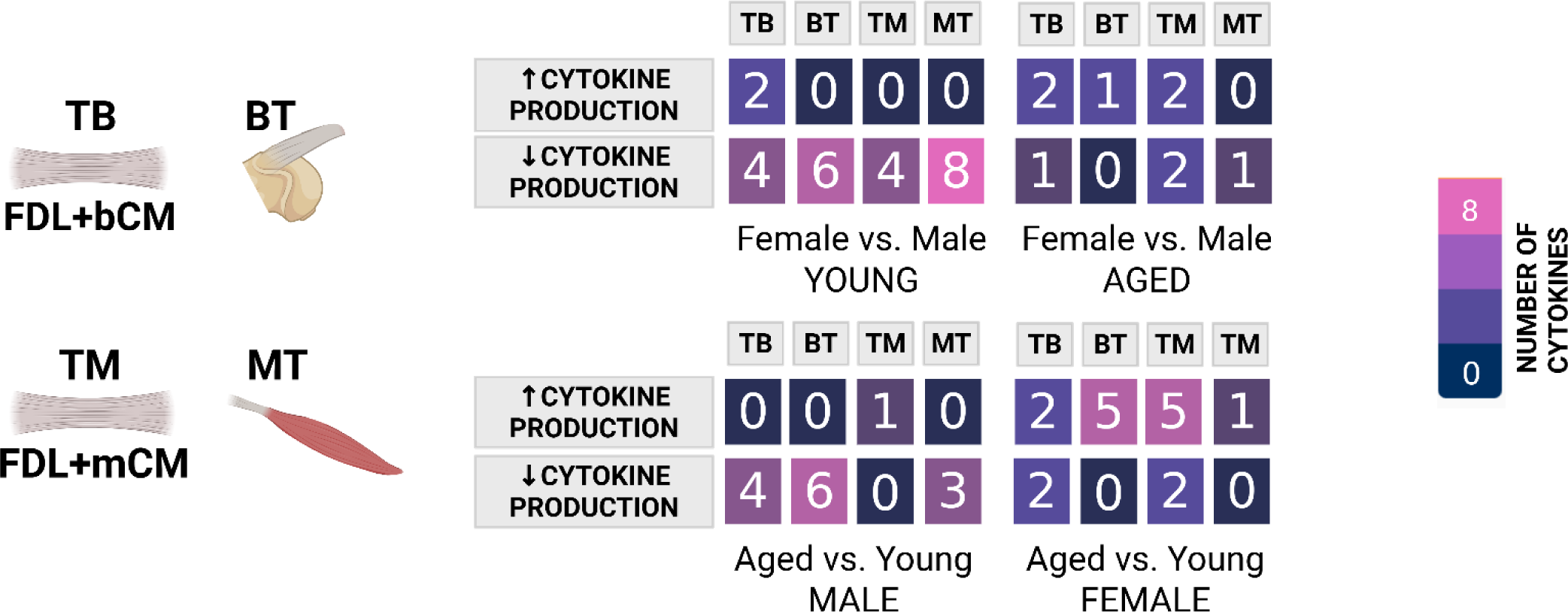
Conditioned media containing secreted factors from injured bone (bCM) or muscle (mCM) were applied to healthy flexor tendon explants for 24 hours to assess the effects of bone and muscle injury on tendon health (TB and TM conditions, respectively). To study direct crosstalk, we included co-culture conditions that preserved native junctions between bone and tendon (BT) or muscle and tendon (MT). Each table displays the number of cytokines that were significantly increased or decreased (normalized to the corresponding tissue-only controls; see Figure 2 caption for details). Data are stratified by age and sex. The color gradient reflects the number of cytokines altered, with dark blue indicating no changes (0 cytokines) and pink indicating up to eight cytokines altered. The exact number of cytokines altered is also noted within each group and comparison.

Aging itself, however, did significantly disrupt tissue signaling, but this was dependent on both sex and the direction/initiator of crosstalk molecules (Fig 3,4,6). Intertissue communication mostly increased with age in females but decreased with age in males. This is unexpected, given previous findings suggesting worse healing outcomes and diminished remodeling responses in females. Given the established role of sex hormones in musculoskeletal health, these results suggest that intertissue communication may contribute to sex-specific musculoskeletal declines with aging. Future studies should further explore the impact of hormonal regulation on these interactions to better understand their implications. Female samples also exhibited significant age-related changes in crosstalk from muscle to tendon, but not from tendon to muscle. Sarcopenia, the aging-associated loss of skeletal muscle mass, quality, and function, has been shown to be affected by sex hormones.^17^ Therefore, it’s possible that these findings are directly related to an aging-associated and sex-dependent phenotype in muscle rather than tendon.

Looking across all conditions (Fig. 6), it appears that bone-tendon crosstalk was more affected by aging than muscle-tendon interactions. This could be due to direct bone-tendon interaction in the synovial space, where secreted proteins dominate crosstalk mechanisms. There have been many studies demonstrating significant overlap in signaling between tendon and bone progenitor populations recently as well, so it’s possible that there are established signaling relationships at the enthesis.^18,16^ It’s not clear why this wouldn’t be similar at the myotendinous junction, although its development has been studied far less.^20^ Overall, we saw fewer changes in muscle to tendon crosstalk with aging, particularly in males (Fig 4b). However, we did find a very strong decrease in secreted IL-15 and IL-6 levels in old muscles with tendon included. Both cytokines are known exercise- and injury-induced myokines, so it’s possible that physical connections are key to crosstalk mechanisms between muscle and tendon.^21,19^

While our *in vitro* co-culture model enables controlled investigation of local crosstalk between tendon, muscle, and bone tissues following injury, it is inherently limited by the absence of systemic interactions present *in vivo*. Without immune cell infiltration, our system cannot fully recapitulate the dynamic and temporally regulated immune responses that occur in living organisms. We are actively developing novel model systems to study this process *in vivo* to overcome these limitations. In the meantime, it’s important to recognize that there is no clearance *in vitro* and therefore cytokine concentrations remain elevated and constant between media changes, potentially leading to amplified or prolonged signaling responses that exceed physiologic levels. We are currently exploring time-dependent collection of medium to investigate secreted factor dynamics. Additionally, the use of fetal bovine serum introduces undefined levels of growth factors, chemokines and cytokines. While FBS levels are consistent across all experiments, lot-to-lot variability may further influence the observed responses. Despite these limitations, our model allows for a unique platform to dissect local tissue-tissue interactions that would be difficult to isolate *in vivo*.

Future studies should aim to further elucidate the dynamic nature of tissue crosstalk in the musculoskeletal system, particularly as it relates to age and sex differences. Temporal profiling of cytokine secretion following injury in co-culture systems would help clarify whether aging alters not only the magnitude but also the timing of signaling responses. Moreover, incorporating mechanical loading into co-culture systems would enable investigation of how physiological forces influence crosstalk, particularly mechano-responsive pathways like IL-6 and IGF-1, which may be altered with age. All three tissues are highly responsive to mechanical loading, which plays a key role in maintaining their structure and function.^23-25^ Although this timeframe is not expected to result in large changes based on mechanical unloading, we will explore this in the future.^25^ Single-cell transcriptomics of co-cultured tissues could also reveal subpopulation-specific signaling networks, including the roles of progenitor cells, immune cells and senescent cell types, and how these populations contribute to altered crosstalk across tissue interfaces. Finally, given the observed sex-dependent differences in muscle-tendon signaling with age, studies evaluating the influence of sex hormones on these pathways, using hormone supplementation, could uncover hormonal regulation of key cytokines and growth factors involved in inter-tissue communication. Together, these approaches would provide a more comprehensive understanding of the biological and biomechanical factors that shape tissue crosstalk with aging.

In summary, our study highlights the complex and dynamic nature of inflammatory crosstalk between muscle, bone, and tendon, emphasizing the influence of age and sex on these interactions. Crosstalk between tissues was highly affected by both aging and sex. Interestingly, secreted interactions appeared to dominate crosstalk between bone and tendon while physical interactions were necessary for muscle-tendon communication. While it’s not clear what role inter-tissue signaling plays in joint health overall yet, we suspect that tissues coordinate responses to exercise and injury to aid in recovery. The disruption to crosstalk in aging could be partially responsible for the increased susceptibility of aging individuals to total joint degeneration, poor healing capacity, and limited exercise responses. The pronounced effect of tendon on inter-tissue inflammatory responses suggests that tendon dysfunction could exacerbate aging-related musculoskeletal deterioration, highlighting tendon as a critical but often overlooked player age-related joint pathology.

## Acknowledgments

This work was supported by NIH R35-GM151127 and the Evans Center Musculoskeletal Health (MHet) Affinity Research Center (ARC).

## Supplementary Figures

**Supplemental Figure 1:**
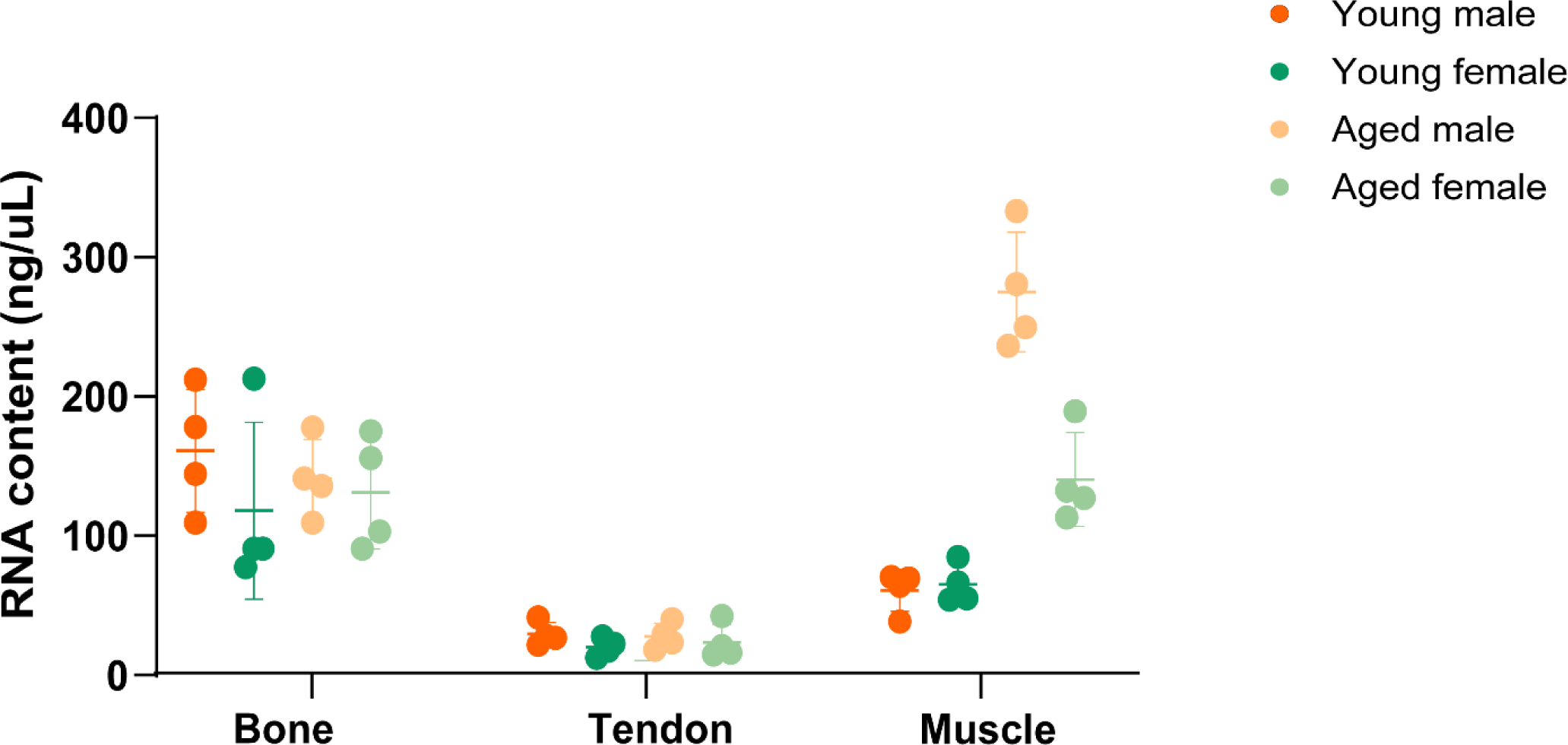
Size comparison of bone, tendon, and muscle explants based on total RNA content. Total RNA was extracted from isolated bone, tendon, and muscle tissues to assess relative tissue size and cellularity. RNA yield (ng/uL) was used as a proxy for tissue mass and cell density. Data are presented as mean ± SD.

**Supplemental Figure 2:**
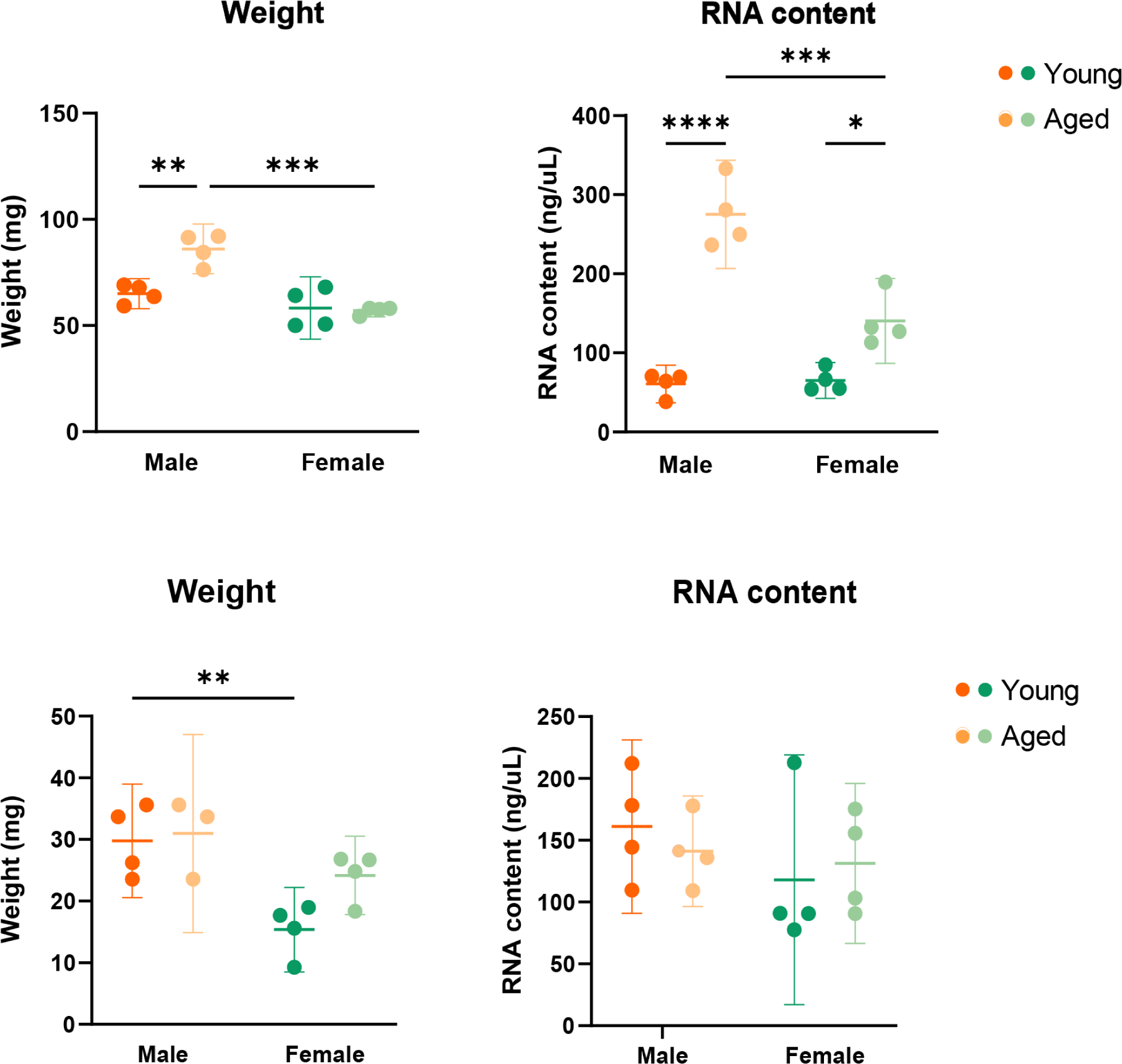
Comparison of weight and RNA content between bone and muscle explants. Freshly isolated bone and muscle tissues were weighed prior to RNA extraction to evaluate differences in tissue mass and cellular content. (a) Tissue weight (mg) and (b) total RNA yield (ng/uL) were quantified for each tissue type. Data are presented as mean ± SD

**Supplemental Figure 3:**
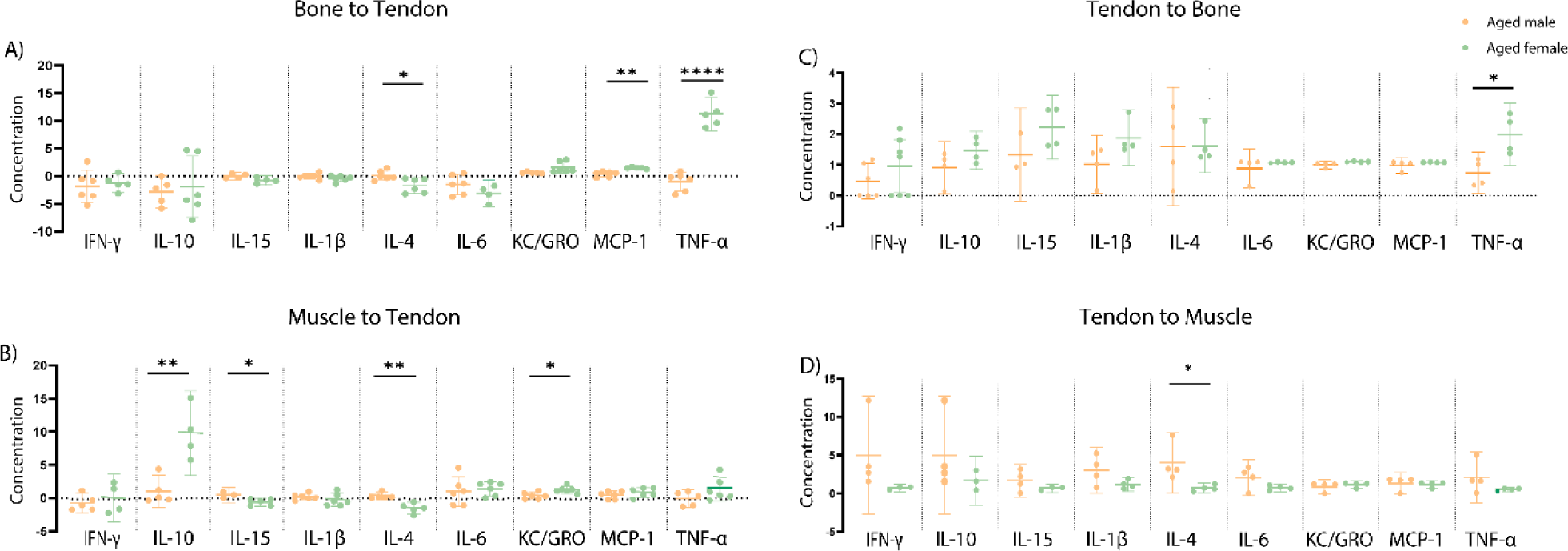
Sex Differences in crosstalk in aged rotator cuff tissues. Concentration of pro- and anti- inflammatory cytokines released in spent media from aged male and female CM treated FDLs (A and B) and BT and MT coculture (C and D). All data points are normalized to bone/tendon/muscle alone. Data shown as mean ± SD. Statistical significance (*p≤0.05) between normalized aged male and female represented by solid line spanning between groups.

## Notes

### Competing Interest Statement

The authors have declared no competing interest.

## References

1. Farley KX, Wilson JM, Kumar A, et al. Prevalence of Shoulder Arthroplasty in the United States and the Increasing Burden of Revision Shoulder Arthroplasty. JB JS Open Access. 2021;6(3):e20.00156. doi:10.2106/JBJS.OA.20.00156

2. Hügle T, Geurts J, Nüesch C, Müller-Gerbl M, Valderrabano V. Aging and osteoarthritis: an inevitable encounter? J Aging Res. 2012;2012:950192. doi:10.1155/2012/950192

3. Baechle JJ, Chen N, Makhijani P, Winer S, Furman D, Winer DA. Chronic inflammation and the hallmarks of aging. Mol Metab. 2023;74:101755. doi:10.1016/j.molmet.2023.101755

4. Lavagnino M, Wall ME, Little D, Banes AJ, Guilak F, Arnoczky SP. Tendon Mechanobiology: Current Knowledge and Future Research Opportunities. J Orthop Res Off Publ Orthop Res Soc. 2015;33(6):813–822. doi:10.1002/jor.22871

5. Arvind V, Huang AH. Reparative and Maladaptive Inflammation in Tendon Healing. Front Bioeng Biotechnol. 2021;9:719047. doi:10.3389/fbioe.2021.719047

6. Connizzo BK, Grodzinsky AJ. Release of pro-inflammatory cytokines from muscle and bone causes tenocyte death in a novel rotator cuff in vitro explant culture model. Connect Tissue Res. 2018;59(5):423–436. doi:10.1080/03008207.2018.1439486

7. Bone and muscle crosstalk in ageing and disease | Nature Reviews Endocrinology. Accessed May 4, 2025. https://www.nature.com/articles/s41574-025-01088-x

8. Perruccio AV, Badley EM, Power JD, et al. Sex differences in the relationship between individual systemic markers of inflammation and pain in knee osteoarthritis. Osteoarthr Cartil Open. 2019;1(1-2):100004. doi:10.1016/j.ocarto.2019.100004

9. Connizzo BK, Grodzinsky AJ. Release of Pro-Inflammatory Cytokines from Muscle and Bone Causes Tenocyte Death in a Novel Rotator Cuff In Vitro Explant Culture Model. Connect Tissue Res. 2018;59(5):423–436. doi:10.1080/03008207.2018.1439486

10. Mihailova A. Interleukin 6 Concentration in Synovial Fluid of Patients with Inflammatory and Degenerative Arthritis. Curr Rheumatol Rev. 2022;18(3):230–233. doi:10.2174/1874471015666220128113319

11. Øvlisen K, Kristensen AT, Jensen AL, Tranholm M. IL-1β, IL-6, KC and MCP-1 are elevated in synovial fluid from haemophilic mice with experimentally induced haemarthrosis. Haemophilia. 2009;15(3):802–810. doi:10.1111/j.1365-2516.2008.01973.x

12. Maruyama M, Rhee C, Utsunomiya T, et al. Modulation of the Inflammatory Response and Bone Healing. Front Endocrinol. 2020;11:386. doi:10.3389/fendo.2020.00386

13. McCarthy M, Raval AP. The peri-menopause in a woman’s life: a systemic inflammatory phase that enables later neurodegenerative disease. J Neuroinflammation. 2020;17(1):317. doi:10.1186/s12974-020-01998-9

14. Gilmer G, Hettinger ZR, Tuakli-Wosornu Y, et al. Female aging: when translational models don’t translate. Nat Aging. 2023;3(12):1500–1508. doi:10.1038/s43587-023-00509-8

15. Klein SL, Flanagan KL. Sex differences in immune responses. Nat Rev Immunol. 2016;16(10):626–638. doi:10.1038/nri.2016.90

16. Qi S, Ngwa C, Morales Scheihing DA, et al. Sex differences in the immune response to acute COVID-19 respiratory tract infection. Biol Sex Differ. 2021;12(1):66. doi:10.1186/s13293-021-00410-2

17. Messier V, Rabasa-Lhoret R, Barbat-Artigas S, Elisha B, Karelis AD, Aubertin-Leheudre M. Menopause and sarcopenia: A potential role for sex hormones. Maturitas. 2011;68(4):331–336. doi:10.1016/j.maturitas.2011.01.014

18. Schwartz AG, Galatz LM, Thomopoulos S. Enthesis regeneration: a role for Gli1+ progenitor cells. Dev Camb Engl. 2017;144(7):1159–1164. doi:10.1242/dev.139303

19. Killian ML. Growth and mechanobiology of the tendon-bone enthesis. Semin Cell Dev Biol. 2022;123:64–73. doi:10.1016/j.semcdb.2021.07.015

20. Charvet B, Ruggiero F, Le Guellec D. The development of the myotendinous junction. A review. Muscles Ligaments Tendons J. 2012;2(2):53–63.

21. Pedersen BK, Steensberg A, Fischer C, et al. Searching for the exercise factor: is IL-6 a candidate? J Muscle Res Cell Motil. 2003;24(2-3):113–119. doi:10.1023/a:1026070911202

22. Wong W, Crane ED, Kuo Y, Kim A, Crane JD. The exercise cytokine interleukin-15 rescues slow wound healing in aged mice. J Biol Chem. 2019;294(52):20024–20038. doi:10.1074/jbc.RA119.010740

23. Wang L, You X, Zhang L, Zhang C, Zou W. Mechanical regulation of bone remodeling. Bone Res. 2022;10(1):1–15. doi:10.1038/s41413-022-00190-4

24. Aggouras AN, Stowe EJ, Mlawer SJ, Connizzo BK. Aged Tendons Exhibit Altered Mechanisms of Strain-Dependent Extracellular Matrix Remodeling. J Biomech Eng. 2024;146(7):071009. doi:10.1115/1.4065270

25. Zhang J, Wang JHC. The Effects of Mechanical Loading on Tendons - An In Vivo and In Vitro Model Study. PLoS ONE. 2013;8(8):e71740. doi:10.1371/journal.pone.0071740

